# Effects of ancient anthropogenic clam gardens on the growth, survival, and transcriptome of Pacific littleneck clams (*Leukoma staminea*)

**DOI:** 10.1101/2022.09.09.507365

**Authors:** Monique R. Raap, Helen J. Gurney-Smith, Sarah E. Dudas, Christopher M. Pearce, Jong S. Leong, Ben J. G. Sutherland, Ben F. Koop

## Abstract

Clam gardens traditionally established and maintained by coastal Indigenous Peoples of northwest North America are habitat modifications to enhance intertidal clam productivity for reliable local food production. In this study, phenotypic and transcriptomic responses of Pacific littleneck clams (*Leukoma staminea*) were evaluated 16 weeks after transplantation to either unmaintained clam gardens or reference (unmodified) clam beaches. Beach sediment characteristics including grain size and organic content were examined across all beaches. Large differences in abiotic characteristics and phenotypic responses were observed among beaches; however, differences were not related to the clam garden/reference beach effect. Clam survival and growth were negatively associated with small rocks, very fine sand, and silt, along with carbonate and organic content, and positively associated with coarse sand, sand, and fine sand. To investigate molecular responses to unmaintained clam gardens, a *de novo* transcriptome containing 52,000 putative transcripts was assembled for *L. staminea* and was used to test for differential expression between transplanted clams in unmaintained clam gardens or on reference beaches. As expected, given the lack of significant phenotypic differences between treatments, transcriptomic responses to unmaintained clam gardens were minor, although several weakly-associated transcripts were identified. By contrast, the strong survival gradient across beaches was used to identify genes associated with survival and, combined with characterization of tissue-specific expression in the gill and digestive gland, contributes to our understanding of molecular processes in this non-model species.

## Introduction

Coastal shorelines have been altered to increase productivity and facilitate the harvesting of natural resources for centuries (Erlandson et al., 2008). Shoreline constructions such as clam gardens and fishing weirs have long been used to provide productive and predictable local food resources for coastal Indigenous Peoples (Neudorf et al., 2017) and provide important insights into cultural food practices, traditional technologies, economies, values, and ancestral practices of coastal communities (Deur et al., 2015; Jackley et al., 2016; Lepofsky et al., 2020; Smith et al., 2019). Construction of clam gardens in North America is estimated to have begun approximately 3,500 years ago (Smith et al., 2019) and clam gardens are found from Alaska (USA), through British Columbia (BC, Canada), to Washington State (USA) (Groesbeck et al., 2014).

Construction of clam gardens involves rolling rocks and boulders to the low-tide line and building a wall parallel to the shoreline (Deur et al., 2015), allowing sediment deposition between the wall and the high-tide mark. This creates a wider intertidal shellfish habitat area with a reduced slope (Neudorf et al., 2017) where shellfish such as Pacific littleneck clams (*Leukoma staminea*) typically occur (Deur et al., 2015; Groesbeck et al., 2014). The reduced slope allows for a thin layer of seawater to be retained on the accumulated sediment, reducing desiccation risk, keeping clams in shallow accessible portions of the intertidal zone, and maximizing submersion time for feeding and therefore growth (Deur et al., 2015). Furthermore, the exterior, ocean-facing side of the wall creates rocky reef habitat for a wide variety of other marine invertebrates, many of which can also be harvested for consumption (Caldwell et al., 2012; Deur et al., 2015; Lepofsky et al., 2017). Traditionally, clam gardens were maintained and tended with practices that included predator exclusion, selective harvesting, and the addition of gravel and crushed shell or shell hash (Deur et al., 2015). The latter may provide settlement cues to larval shellfish as well as protect young recruits from seawater acidity and predators (Green et al., 2012; Sponaugle & Lawton, 1990).

Research on the effects of clam gardens is an area of active interest. Many clam garden walls, although maintained into the 20^th^ century (Williams, 2006), now remain as unmaintained structures, which could be expected to provide only partial effects relative to a fully tended clam garden. Nonetheless, untended clam gardens have been found to harbour higher densities of Pacific littleneck clams and butter clams (*Saxidomus giganteus*), to have increased recruit survival, and higher growth rates relative to unmodified beaches (Groesbeck et al., 2014; Jackley et al., 2016). Clam gardens also support distinct biological communities with increased abundances of other infaunal taxa including Nematoda, Harpacticoida, and Chironomidae (Cox et al., 2019), and epifaunal taxa such as *Chthamalus dalli*, *Lottia persona*, *Balanus crenatus*, and *Littorina scutulata*, (Cox et al., 2024a). These increases in diversity and density were positively correlated with the quantity of gravel and shell hash present (Cox et al., 2024a; Cox et al., 2019). Sediment composition can be altered in clam gardens, for example through increased carbonate and reduced silt (Salter, 2018), as well as more gravel and shell hash (Groesbeck et al., 2014) relative to unmodified beaches, which are often primarily composed of silt, sand, and mud (Jackley et al., 2016). Such differences in sediment composition are attributed to the tending practices of Indigenous Peoples, which included the addition of gravel and crushed whole clam shells to clam gardens (Groesbeck et al., 2014; Williams, 2006).

The effect of sediment shell hash on clam productivity is not definitive. Sediment carbonate saturation state can affect settlement and recruitment of larval and juvenile bivalves, respectively (Green et al., 2012). Seawater calcium carbonate (CaCO_3_) saturation state is important to the formation and maintenance of shell thickness and integrity of marine molluscs and other calcifiers (Chadwick et al., 2019; Evans et al., 2014; Green et al., 2012; Waldbusser & Salisbury, 2014). Increased sediment shell hash has been observed to increase clam settlement and productivity (Green et al., 2012; Groesbeck et al., 2014; Salter, 2018), but observations have also been made showing negative (Munroe, 2016) or neutral (Greiner et al., 2018) effects on early post-settlement bivalve growth and recruitment.

Genomic tools such as transcriptomics can capture early signals of environmental stressors and are increasingly used to evaluate abiotic impacts on aquatic species (Milan et al., 2011; Sutherland et al., 2012). Transcriptomic data can be associated with phenotypic (*e.g.*, survival, growth) and environmental data and used to predict potential physiological outcomes (Evans et al., 2011; Miller et al., 2017). An additional benefit is that in some studies, metatranscriptomics can provide additional taxonomic identification of microbiota in samples (Bourlat et al., 2013; Sutherland et al., 2022). The present study aims to determine whether Pacific littleneck clam health and productivity—as assessed by growth, survival, and transcriptomic response—are affected when clams are transplanted to unmaintained clam gardens relative to unmodified clam beaches, and whether these unmaintained clam gardens contain significantly different abiotic conditions. To this end, we generated and assembled a *de novo* reference transcriptome and characterized Pacific littleneck clam transcriptome responses in several unmaintained clam gardens and unmodified beaches in the same area. Beaches used in the study were in northern Quadra Island in coastal BC, an area with a very high density of historic clam gardens, covering approximately 35% of the shoreline (Groesbeck et al., 2014; Lepofsky et al., 2020; Neudorf et al., 2017). To evaluate the effects of the unmaintained clam gardens on transplanted clams after a 16-week growth period, measured environmental parameters, clam phenotypic responses (growth and survival), and clam transcriptomic responses were considered. The combination of abiotic measurements and organismal and molecular phenotypes with the characterized transcriptome provides valuable context for both the effects of different environmental variables on Pacific littleneck clams and on the functions of response genes in this non-model, ecologically and culturally important species.

## Methods

### Positionality Statement

The authors of this paper are academic, government, and independent scientific researchers who conduct research on climate change, ecology, invertebrate physiology, bioinformatics, and genomics. The authors were educated and trained in North American and European institutions prior to the positions in Canada that they hold at time of submission, with five reaching doctorate status and two with master’s degrees. This research was conducted in British Columbia, Canada. None of the authors who have provided heritage information for this statement indicated First Nation or other Indigenous background, and therefore we respectfully acknowledge the potential for unconscious bias on the presented subject matter.

### 2.1 | Study sites

The intertidal coastlines of Kanish Bay and adjoining Small Inlet on northwest Quadra Island, BC (Figure 1), were chosen for this study due to an abundance of historic clam gardens where sea level has remained relatively consistent through time, ensuring walls were still at an ecologically optimal height, and due to the proximity to the Hakai Institute research station that supported field logistics. Also, complimentary research has been conducted on clam gardens in the same region and on many of the same sites (Cox et al., 2024a; Cox et al., 2024b; Groesbeck et al., 2014; Lepofsky et al., 2020).

**FIGURE 1.**
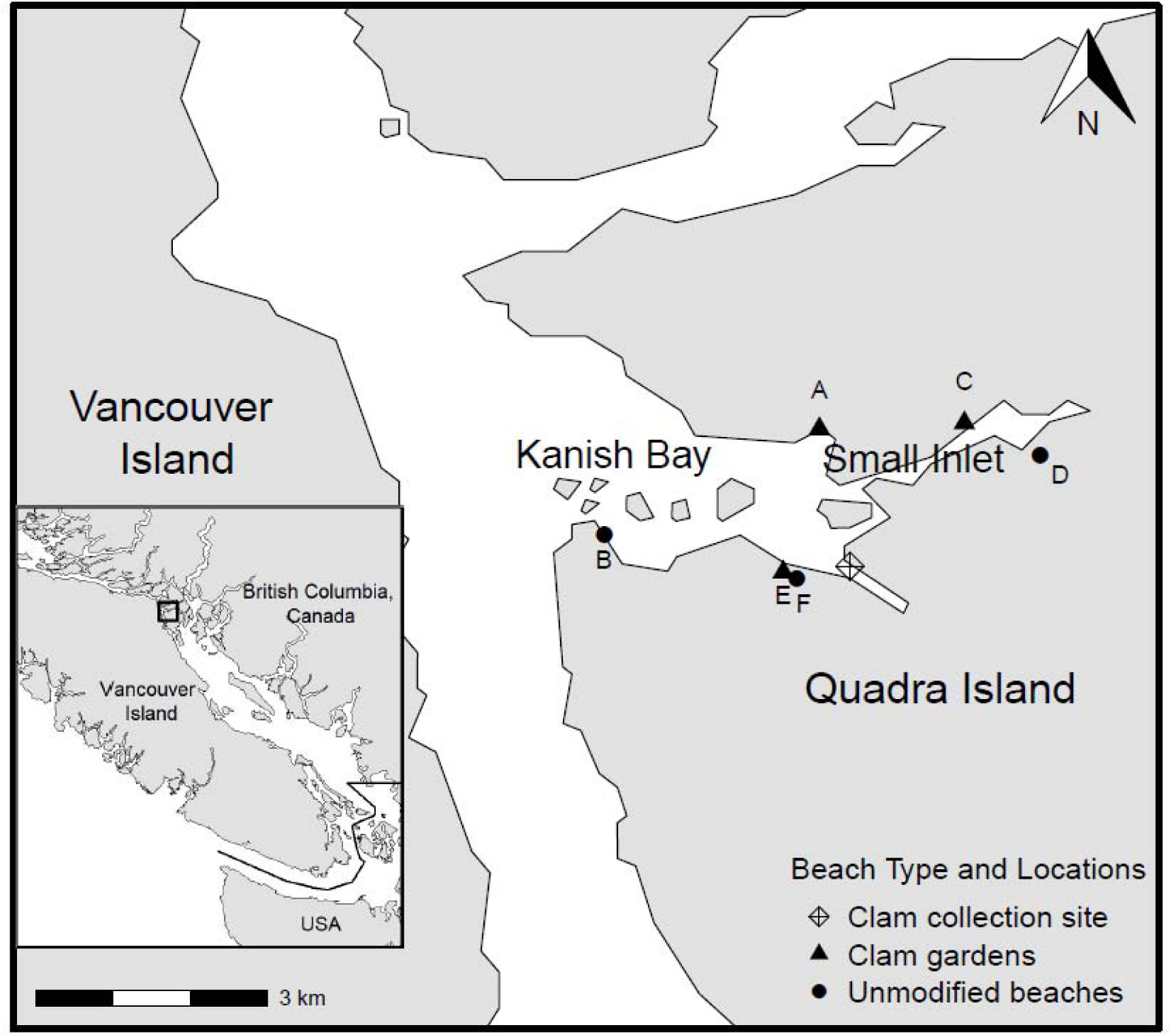
Locations of unmaintained clam gardens (A, C, and E; triangles) and unmodified reference clam beaches (B, D, and F; circles) used in the study. The collection site used as source for the clams used in the study is designated by a diamond. Inset map shows the location of the study site in western Canada.

Quadra Island is located off the northeast coast of Vancouver Island and is in the traditional territories of the Kwakwaka’wakw (Laich-kwil-tach) and northern Coast Salish Peoples (Smith et al., 2019). Three unmaintained clam garden beaches (CG, designated as A, C, and E) were selected for the study along with three unmodified reference clam beaches (Ref, designated as B, D, and F) (Figure 1). Beaches with similar exposure, slope, and sediment (all determined in the field visually) were sought to reduce the influence of other variables as much as possible. Once three reference sites were identified, nearby clam gardens with intact walls were selected. Additionally, all sites were pre-surveyed for the presence of live bivalve populations using test digs. After the study began, beach B was identified as a damaged clam garden with over 90% of its wall destroyed most likely due to logging activities in the early 1900s that involved dragging log bundles down the beach into the ocean (Twindle, 1918). Now only a small remnant of the wall is visible at the very edges at tidal heights below Mean Lower Low Water (MLLW). The clam gardens are not believed to have been tended as per Indigenous practices since the early-to-mid 1900s (Deur et al., 2015).

### 2.2 | Clam transplantation experiment

Juvenile Pacific littleneck clams (N = 400) of 1−2 cm in shell height (hinge to shell margin) were hand dug from a site in Kanish Bay (Figure 1) between 1 and 2 m MLLW level on May 7, 2016. Each clam was individually wet weighed, measured for shell height and length (*i.e.*, the widest part of the shell at 90° to the shell height), and then haphazardly placed into 18 groups of 20 clams each. Clams were held in a pearl net below the low intertidal zone in Heriot Bay (Quadra Island) until deployment. Clams on beach A were deployed on May 10, those on beaches B, C, and D on May 11, and those on beaches E and F on May 12, 2016, different deployment days being required due to tidal heights and weather conditions. At low tide at each of the six beaches, the 1.5–1.8 m intertidal zone was demarcated and three plastic mesh cubes (length x diameter x height: 20 x 20 x 4 cm; made from black high-density polyethylene, Vexar^™^), each containing 20 clams, were buried in the top 20 cm of the sediment, 5 m apart and parallel to the shore in the middle of the demarcated zone. Each mesh cube was defined as one replicate plot (P) and uniquely numbered: Beach A with P1–P3, Beach B with P10−P12, Beach C with P4−P6, Beach D with P7−P9, Beach E with P13–P15; and Beach F with P16–P18.

Transplanted clams were left *in situ* for 16 weeks and collected on August 30–31, 2016. Each mesh cube, still containing both sediment and clams, was carefully removed from the beach with minimal disturbance. The cubes were then individually placed in bags and transported in coolers to the Hakai Institute laboratory on Quadra Island. Upon arrival at the laboratory, each individual was processed as quickly and carefully as possible. Immediately after wet weight, shell height, and shell length were measured, and the status (alive or dead) recorded, tissue samples were collected for gene expression analysis.

Growth and survival per plot were calculated as:

*Percent growth* = (mean final height – mean initial height) / mean initial height x 100
*Percent survival* = number of survivors / initial number of clams x 100

In addition, some visual observations were made on animal condition, following the protocols developed for the Canadian National Animal Aquatic Health Program (NAAHP), including observations of animal state (*i.e.*, health/vitality, condition) and digestive gland colouration.

### 2.3 | Sediment collection and analysis

Sediment-core samples (length x diameter: 15 x 5 cm) were collected adjacent to each deployed cube (*n* = 3 cores per beach) to assess grain size. Additional sediment-core samples (10 x 3.5 cm) were collected adjacent to each cube to assess percent carbonate and percent organic matter content (*n* = 3). Each core was placed individually in labeled sample bags and frozen at −20^°^C until analysis.

To determine grain-size distribution, the sediment samples were dried to constant weight at 100^°^C for 24 hr, weighed, transferred to a series of nested sieves (mesh size: 4.75 mm, 2 mm, 1 mm, 500 µm, 250 µm, 125 µm, and 63 µm), and shaken in a sediment shaker for 15 min. Each size fraction was then weighed and expressed as a percentage of the total sediment weight. A scale very similar to the Wentworth scale of sediment size classes was used to classify the grain sizes: pebble and larger gravel called “rocks” in the present study (> 4.75 mm), granule gravel called “small rocks” in the present study (2.00 – 4.75 mm), very coarse sand (1.00 – 2.00 mm), coarse sand (0.50 – 1.00 mm), sand (250 – 500 µm), fine sand (125 – 250 µm), very fine sand (63 – 125 µm), and silt (< 63 µm) (Wentworth, 1922).

The loss on ignition method (Heiri et al., 2001) was used to determine sediment percent organic matter and carbonate content. Briefly, the sediment samples were dried to constant weight at 100^°^C for 48 hr in previously-ashed (450^°^C for 8 hr) crucibles, and the dry weights measured. Dried sediment samples were then ashed in a muffle furnace at 435^°^C for 8 hr. After cooling the samples in a desiccator for 1 hr, the weights of the ashed samples were recorded. The percent organic matter was then calculated using the following equation:

*Percent organic matter* = (dry weight – ash weight) / dry weight x 100

Sediment percent carbonate content was determined following 2 hr at 950^°^C and 2 hr of cooling in a desiccator, and calculated using the equation:

*Percent carbonate content* = (ash weight_435_ – ash weight_950_) / ash weight_435_ x 100

### 2.4 | Data analysis: clam size, growth and survival, and sediment characteristics

Data were analyzed in R v.4.4.2. (R.Core.Team, 2025). Model assumptions of normality were assessed using histograms, quantile-quantile plots of the data, and the Shapiro-Wilk test. Homogeneity of the residuals was assessed using plots of model residuals versus fitted values and Bartlett’s test for homogeneity of variance. Nested ANOVAs (plots nested within beach location), using the linear mixed-effects function in the *nlme* package, were used to test for differences between clam garden and unmodified reference beaches (*i.e.*, beach type, two levels) in initial and final clam shell heights, wet weights, percent growth, and percent survival as well as individual sediment characteristics (*i.e.*, rocks, small rocks, very coarse sand, coarse sand, sand, fine sand, very fine sand, silt, percent organic matter, and percent carbonate concentration), with beach location (six levels) included as a random factor. Significance was considered when *p* ≤ 0.05. ANOVAs, linear mixed-effects models, and post-hoc least squares means omitting beach type were then used to test for significant differences among beaches in percent growth, percent survival, sediment carbonates, organics, and individual grain sizes. Principal component analysis (PCA) and biplot figures were generated in R using the prcomp function in the *stats* package and the *ggbiplot* function/package. Pearson correlation between the variables was evaluated with the cor function of the *stats* package of R and inspected with *corrplot* v.0.92 (Wei & Simko, 2021). All code for the analysis is available (see *Data Availability*).

### 2.5 | RNA sampling and extraction

For gene-expression analysis, small sections (∼2 x 2 mm) of gill and digestive gland tissues were excised from each surviving clam using sterile techniques and stored in RNAlater as per manufacturer’s (Ambion) protocols. These two tissues were selected because the bivalve gill is often used to study rapid environmental responses and the digestive gland is an accumulatory organ often used in toxicological and immunology studies for longer-term effects of environmental change and exposure (Gosling, 2008; Milan et al., 2011). Total RNA from tissue sections of 25–30 mg was individually extracted from each tissue by homogenization in 2-mL tubes containing Lysing Matrix D (MP Biomedicals) in a Tissuelyser II (Qiagen) at 25 Hz for 2 min followed by purification through RNeasy columns (Qiagen). To eliminate DNA contamination, a DNase protocol was applied (Turbo DNA-free Kits, Ambion). After DNase treatment, total RNA was quantified by spectrophotometry (NanoDrop 1000, Thermo Fisher Scientific).

### 2.6 | Library preparation for RNA sequencing

Up to three pooled samples per beach were created for each of the six beaches by combining five haphazardly selected individuals from each plot within the beach. Plots were only used if they contained at least five surviving clams. This resulted in three plots being used per beach for all locations except beaches E and F, which only had five or more survivors in a single plot per beach. This resulted in 14 RNA pools per tissue (N = 28 pools total) to be used for library synthesis and sequencing. Five total RNA samples were normalized to generate equimolar pools of RNA for each tissue for the 14 plots across the six beaches. Pooled RNA quality and quantity were evaluated using the RNA 6000 Nano chip on a 2100 Bioanalyzer (Agilent).

Libraries were generated using 250 ng of pooled total RNA using an mRNA enrichment with the NEBNext Poly(A) Magnetic Isolation Module (New England Biolabs) and cDNA synthesis was conducted using NEBNext RNA First Strand Synthesis and NEBNext Ultra Directional RNA Second Strand Synthesis Modules. Remaining library preparation used an NEBNext Ultra II DNA Library Prep Kit for Illumina, with adapters and PCR primers (NEB). The remaining steps of library preparation were taken using the Quant-iT^™^ PicoGreen^®^ dsDNA Assay Kit (Life Technologies) and the Kapa Illumina GA with Revised Primers-SYBR Fast Universal Kit (Kapa Biosystems (Pty) Ltd.). Average fragment size was determined using a LabChip GX instrument (PerkinElmer). Libraries were sequenced across four lanes of an Illumina HiSeq4000 PE 100 platform (Illumina) and the mean (± SD) read pairs per library was 54.4 ± 9.5 million. All library preparation and sequencing steps were undertaken at the Genome Québec Innovation Centre (Montreal, Canada).

### 2.7 | Transcriptome assembly and annotation

The 28 sequenced libraries were used to generate a *de novo* transcriptome assembly using Trinity 2.5.1 (--min_kmer_cov 2) (Grabherr et al., 2011). Before assembly, reads were trimmed to remove adapters, low quality bases, and too-short reads using Trimmomatic 0.36 (ILLUMINACLIP:TruSeq3-PE-2.fa:2:30:10 LEADING:3 TRAILING:3 SLIDINGWINDOW: 4:15 MINLEN:36) (Bolger et al., 2014). The TransDecoder 5.0.1 pipeline (Haas et al., 2013) was used to predict likely coding sequences in the resultant assembly using homology (pfam-a, Sept. 2017 release, UniProt) and open reading frame (ORF) information. A minimum cutoff for ORFs was 30 amino acids instead of the default 100 amino acids. The best representative transcript was selected for each gene based on TransDecoder predicted ‘complete’ ORF type, which required protein homology. In cases where there were multiple transcripts with complete ORFs for a gene, the transcript with the largest ORF was chosen. Further filtering was carried out to choose only transcripts ≥ 100 base pairs to remove small transcripts that could cause analysis errors, and scanning pfam-a and Uniprot annotations were used to remove repetitive element-related keywords to produce the final assembly, where each transcript putatively represents a single gene.

### 2.8 | Data analysis: RNA sequencing data

Raw reads were imported to the repository *Simple_reads_to_counts* (see *Data Availability*) for quantitation. Raw reads were gently trimmed for quality and to remove adapters using Trimmomatic (Bolger et al., 2014), with the following flags: ILLUMINACLIP:$VECTORS:2:30:10; SLIDINGWINDOW:20:2; LEADING:2; TRAILING:2; MINLEN:80. Both the raw and the trimmed data were inspected using FastQC (Andrews, 2010) and multiQC (Ewels et al., 2016). The *de novo* reference transcriptome (see above) was indexed using bowtie2 (Langmead & Salzberg, 2012) and reads were aligned against it in local mode, allowing 40 alignments to be retained per read (-k 40). To compare with these results, the Manila clam (*Ruditapes philippinarum*) genome (GCA_009026015.1_ASM902601v1) (Yan et al., 2019) was also tested as a potential reference by indexing the genome using hisat2 (Kim et al., 2015) and aligning the Pacific littleneck clam reads against the Manila clam genome (also retaining a maximum of 40 alignments per read; -k 40). The alignments against the Pacific littleneck clam *de novo* reference transcriptome were quantified using eXpress using default parameters (Roberts & Pachter, 2013). The alignments against the Manila clam genome were also quantified using eXpress default parameters but with the flag *--rf-stranded*. Resultant effective read counts quantified against the *de novo* reference transcriptome were used in downstream analyses.

Transcriptome analyses were all conducted as described in the repository *ms_clam_garden* (see *Data Availability*). Transcript annotation, effective counts, and phenotypic data were all imported into R to be analyzed with *edgeR* v.3.36.0 (Robinson et al., 2010). Separate datasets were constructed including all samples, or including only gill or only digestive gland (DG) samples (*n* = 3 datasets). Each dataset was input into a DGEList, then filtered for low expression by requiring at least 10 reads to be aligned to the transcript in at least five individuals (*i.e.*, counts per million (cpm) > 0.82–1.03, depending on the dataset). Samples were normalized by TMM normalization as applied by *calcNormFactors()* and multidimensional scaling (MDS) plots were produced for each of the three datasets.

Tissue-specific expression was determined by identifying transcripts expressed in one tissue and not in the other. The normalized counts for these transcripts were then obtained from the all-sample dataset and 200 of the highest expressed tissue-specific transcripts from both tissues were plotted using *heatmap()* of the *stats* package in R. Gene Ontology (GO) enrichment was conducted using the tissue-specific transcripts from both tissues compared to all expressed transcripts for that tissue. Enrichment analysis was conducted in DAVID Bioinformatics (Huang et al., 2007).

Differential expression analysis was conducted using *limma* v.3.50.1 (Ritchie et al., 2015). Both tissues were analyzed separately and differentially expressed genes were identified using beach type (*i.e.*, CG vs. Ref) as an explanatory variable using a gene-wise negative binomial generalized linear model with quasi-likelihood tests (glmQLFit) and retaining all transcripts with *p* ≤ 0.001. Differential expression, based on beach-type with a Benjamini-hochberg multiple test correction, was also investigated. Differentially expressed genes with *p* ≤ 0.001 were also identified based on survivorship, where the percent survival of each plot was used to bin the plot into one of high, medium, or low survival based on the fourth quartile, second and third quartiles, and the first quartile of percent survival, respectively. Statistical significance was conducted as above, but using contrasts between each survival group. All other analyses and plots were conducted in R and are available in the repository listed above.

## Results

### 3.1 | Clam growth and survival

There were no significant differences between unmaintained clam garden (CG) and reference (Ref) beaches in any measured growth or survival variable (ANOVAs, *p* ≥ 0.168, Table 1). The mean (± SD) initial clam wet weight and shell height for CG and Ref beaches was 1.5 ± 0.2 g and 1.8 ± 0.3 g and 1.5 ± 0.1 cm and 1.6 ± 0.1 cm (N = 180), respectively (Table 1). After 16 weeks *in situ*, these increased to 3.9 ± 1.3 g and 3.8 ± 2.9 g and 1.8 ± 0.3 cm and 1.9 ± 0.4 cm, which in terms of height, was an increase in growth of 20.5 ± 15.3% and 18.0 ± 22.6%, respectively. Calculating percent growth using shell height includes both the alive and dead clams and is therefore considered to be a more accurate measurement of growth compared to using wet weights. Wet weight only includes surviving clams, and therefore does not capture the reduced growth that may have occurred in clams that eventually died. Clam survival was 63.3 ± 28.3% and 58.3 ± 37.0% in CG and Ref beaches, respectively.

**TABLE 1.**
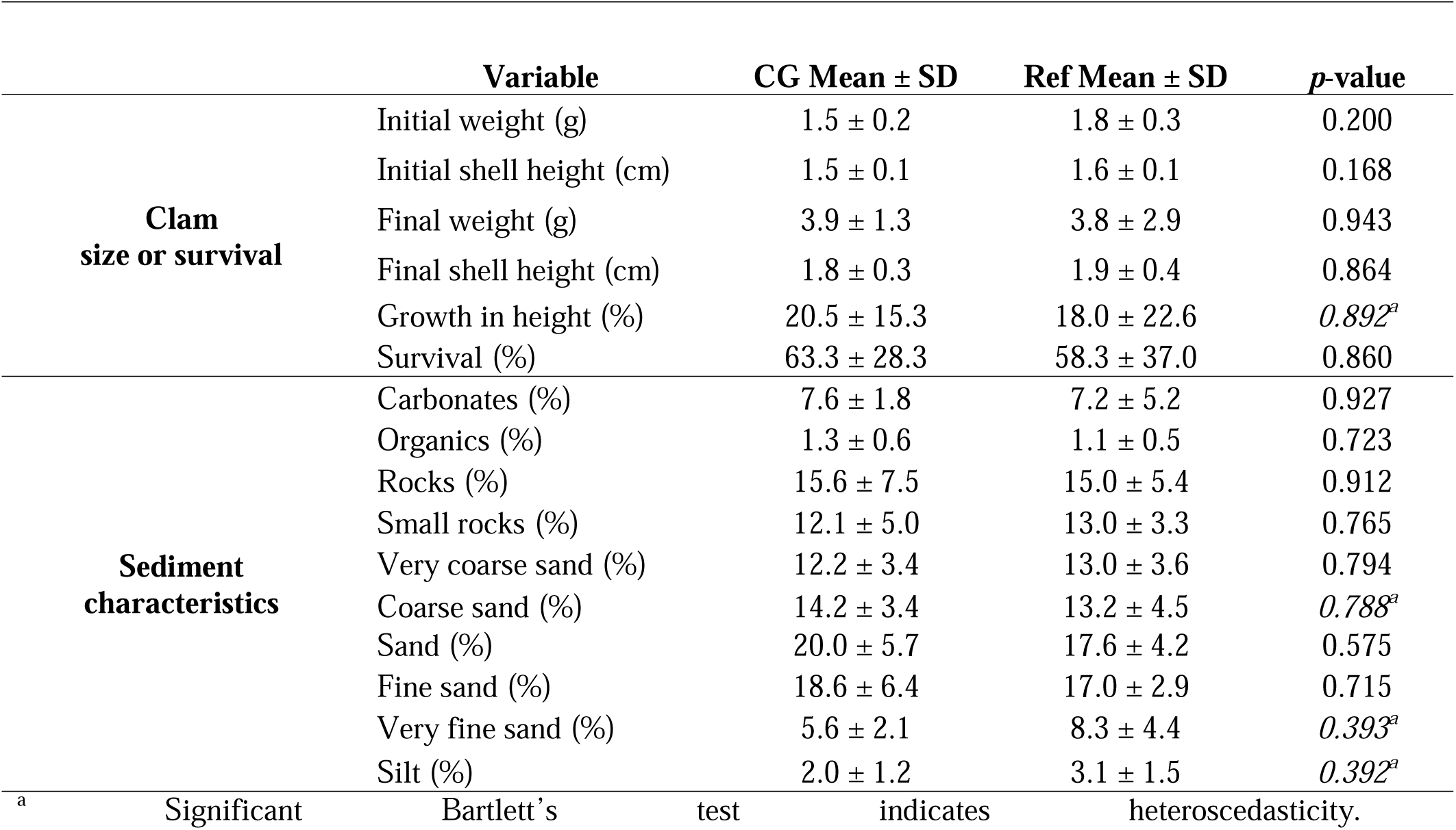
Mean (± SD) clam size or survival and sediment characteristic variables for clam garden (CG) and reference (Ref) beaches and associated linear model *p*-values (with beach location as a nested effect). No variables were significantly affected by the CG/Ref factor. All variables were found to have a normal distribution.

Beach location regardless of beach type significantly affected all growth and survival variables (ANOVAs, *p* ≤ 0.03, Table 2, *SI* Figure S1). An unplanned significant difference occurred in the initial size of clams among the six beaches (initial wet weight and shell height ANOVAs *p* = 0.031 and *p* = 0.044, respectively, Table 2, *SI* Figure S2). This difference was likely due to significantly larger clams inadvertently being planted on beach D (Table 2, *SI* Figure S2). Percent growth was highest on beaches A (31.2 ± 7.2%), C (29.7 ± 2.8%), and D (47.0 ± 8.2%) and lowest on beaches B (6.6 ± 6.8%), E (0.8 ± 0.9%), and F (0.3 ± 0.6%) (Table 2). Percent survival was highest on beaches A (88.3 ± 12.6%), B (71.7 ± 27.5%), C (68.3 ± 12.6%), and D (86.7 ± 5.8%) and lowest on beaches E (33.3 ± 23.6%) and F (16.7 ± 24.7%) (Table 2).

**TABLE 2.**
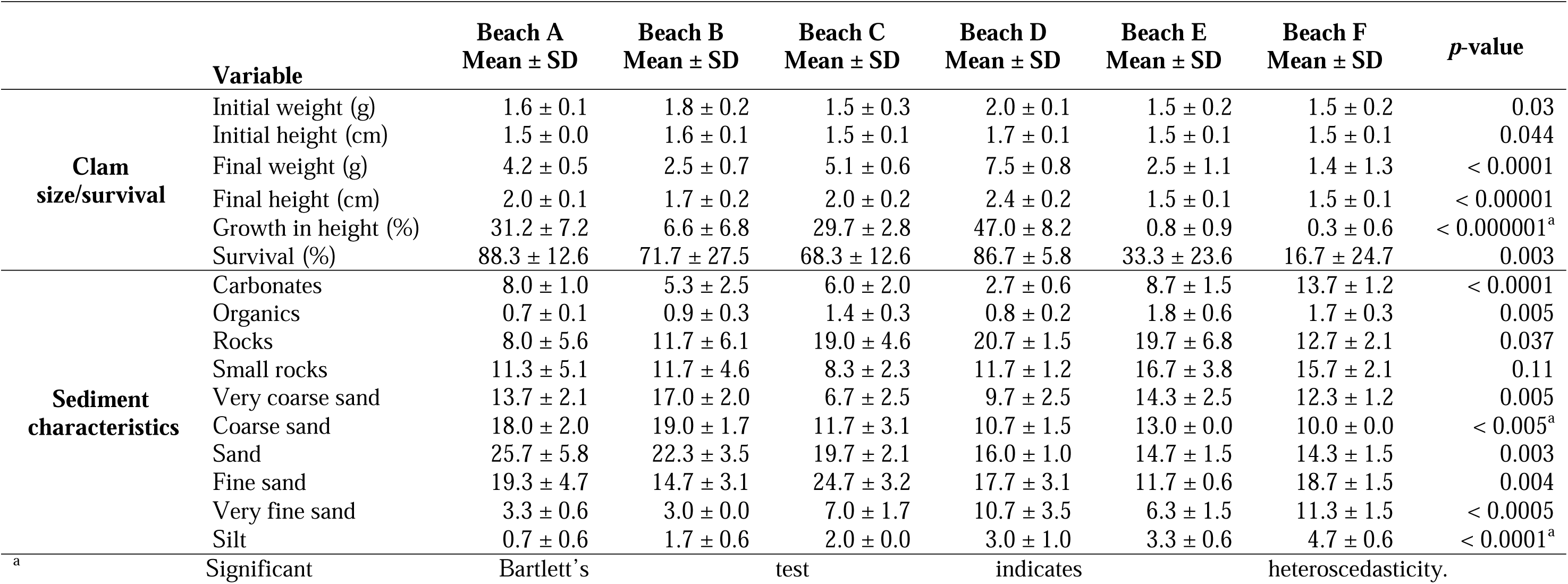
Mean (± SD) clam size/survival and sediment characteristic variables for each beach and associated linear model *p*-values.

Gross observations of the clam showed that most of the survivors on beaches A (CG), B (Ref), C (CG), and D (Ref) were considered healthy (Table 3). These beaches had 68–88% survival, with generally low percentages of clams considered weak (0–5%), emaciated (0–5%), or having pale or very pale digestive glands (0–22%) (beach D was a slight exception, where although survival was high, 22% of clams had pale digestive glands) (Table 3). In contrast, survivors from low-survival (17–33%) beaches E (CG) and F (Ref) had more clams considered weak (13–15%), emaciated (15–22%), and with digestive glands being very pale (13–22%). Beaches E and F are therefore considered poorer performing sites.

**TABLE 3.** Gross observations (health, condition, and digestive gland coloration) of surviving clams in each beach (A−F)

| TMetric | Beach A | Beach B | Beach C | Beach D | Beach E | Beach F |
| --- | --- | --- | --- | --- | --- | --- |
| Health (% weak) | 0 | 5 | 0 | 2 | 13 | 15 |
| Condition (% emaciated) | 0 | 5 | 0 | 2 | 22 | 15 |
| Digestive gland (% pale) | 8 | 7 | 5 | 22 | 0 | 0 |
| Digestive gland (% very pale) | 0 | 0 | 0 | 2 | 22 | 13 |

### 3.2 | Beach sediment characteristics

There were no significant differences between unmaintained CG and Ref beaches for any of the sediment characteristics (ANOVAs, *p* ≥ 0.392, Table 1). However, beach location regardless of beach type significantly affected all sediment characteristics except percentage of small rocks (Table 2). A PCA of sediment characteristics explained 66.0% (dimension 1: 40.8%, dimension 2: 25.6%) of the variation in the data and separated beach locations into three groups (Figure 2), where pairs of beaches were observed to be clustered based on similar geographic locations within Kanish Bay (Figure 1). Beach A (CG), on a west-facing bay, was exposed to wave action from within and outside Kanish Bay. Beach A grouped with beach B (Ref), an east-facing bay that was also exposed to wave action within Kanish Bay (Figure 1).

**FIGURE 2.**
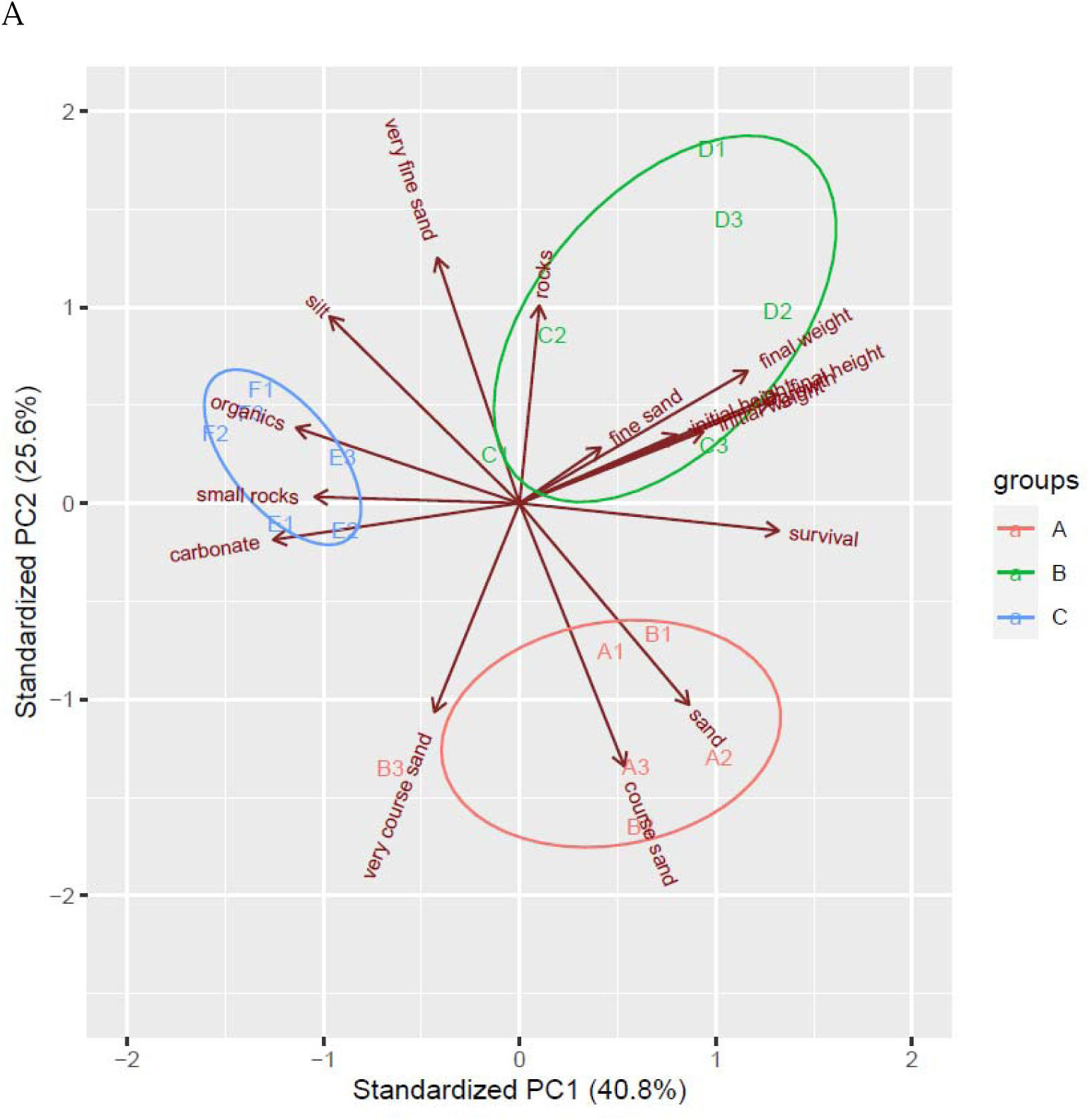
Principal component analysis biplot based on abiotic and biotic phenotypes showing beach groupings and correlations. Each beach is denoted as A through F as per Figure 1 locations, and each replicate plot within the beach as 1–3. Colours of plots and ellipses are for visualization purposes only.

These beaches are the two sites that are closest to, but on opposite sides of, the entrance of Kanish Bay and show correlation with very course sand, coarse sand, and sand (Figure 2). Beach C (CG) is a south-facing sheltered beach within Small Inlet and groups with beach D (Ref), a north-facing beach on the opposite side of Small Inlet. Beaches C and D correlated with rocks, fine sand, and final weight/shell height (Figure 2). Beaches E (CG) and F (Ref), two proximate beaches on the south side of Kanish Bay, face north and are exposed to moderate wave action. These sites were associated with carbonates, organics, and small rocks, and showed an inverse correlation with percent survival (Figure 2). Notably, these beach groupings in the PCA remain constant when only abiotic characteristics are included in the PCA, or when only survival and growth variables are included (*data not shown*). The PCA was also run with all data, but excluding beach D due to the aforementioned initial size difference of this beach, and this showed similar associations and groupings to the above, but beach C was on its own (*SI* Figure S3).

Most sand types, silt, and rocks were either positively or negatively correlated with growth and survival variables (*SI* Figure S4). To investigate correlations between sediment characteristics and clam growth and survival further, beach D was again excluded as described above (*SI* Figure S5). Abiotic variables with significant positive correlations with growth included sand (*p* = 0.018, R^2^ = 0.31) and fine sand (*p* = 0.022, R^2^ = 0.29), with those having negative associations being silt (*p* = 0.002, R^2^ = 0.49) and small rocks (*p* = 0.014, R^2^ = 0.34). Variables with significant positive correlations with survival were coarse sand (*p* = 0.014, R^2^ = 0.34) and sand (*p* = 0.002, R^2^ = 0.48), with those having negative associations being carbonates (*p* = 0.008, R^2^ = 0.39) and organic content (*p* = 0.035, R^2^ = 0.24), as well as small rocks (*p* = 0.021, R^2^ = 0.30), very fine sand (*p* = 0.002, R^2^ = 0.49), and silt (*p* = 0.002, R^2^ =0.51) (*SD* Additional File S1, without Beach D).

### 3.3 | Transcriptomic overview and *de novo* reference transcriptome

The gill and digestive gland (DG) libraries produced on average (± SD) 53.8 ± 10.7 and 55.0 ± 8.3 M read pairs per library, respectively. All 28 libraries, comprising 305 GB of high-quality sequence data, were used to create a *de novo* transcriptome assembly (Table 4; see *Data Availability*). The initial Trinity assembly resulted in 1,695,678 transcripts, of which 1,277,478 were likely coding sequences as predicted by TransDecoder. Following filtering of the *de novo* assembly (see *Methods*), there were 52,000 representative transcripts in the assembly, each putatively representing a single gene, and 42,708 had associated Uniprot identifiers.

**TABLE 4.** Overview of RNA-seq reads and quality scores. Totals that include both reads in the pair are indicated by (R1 + R2). Values are shown in billion (B) or million (M)

| *Leukoma staminea* assembly | Value |
| --- | --- |
| Total input reads (R1 + R2) | 3.0 B |
| Total input nucleotides (R1 + R2) | 305 B |
| Average (± SD) input reads per: | |
| single HiSeq4000 lane (R1 + R2) | 762 ± 13.0 M |
| library (read pairs) | 54.4 ± 9.5 M |
| gill library (read pairs) | 53.8 ± 10.7 M |
| digestive-gland library (read pairs) | 55.0 ± 8.3 M |
| Assembly contig length range | 102 – 29,916 |
| Mean (± SD) Phred quality score | 38.9 ± 0.3 |
| Base-calling accuracy | > 99.9% |

Alignments of trimmed reads resulted in a mean (± SD) alignment rate of 27.87 ± 2.48% against the Pacific littleneck clam *de novo* transcriptome. For comparison, the alignment rate against the Manila clam genome (*i.e.*, the closest relative with a reference genome at the date of the present analysis) resulted in very low alignments of 1.10 ± 0.17% and 2.59 ± 1.00% for gill and DG, respectively. All downstream analyses used the *de novo* transcriptome quantified transcripts. After filtering the all-sample dataset (N = 28 pooled samples), 33,825 transcripts (67.7%) were expressed in at least five samples above the applied threshold (see *Methods*).

An MDS plot of samples from both tissue types showed that gill samples were distinctly separate from DG samples across the first dimension, explaining 65% of the variation in the dataset (Figure 3A). The second dimension, which explained only 3% of the variation, separated the DG samples, but not the gill ones. One outlier sample, P12, was observed in the gill data.

**FIGURE 3.**
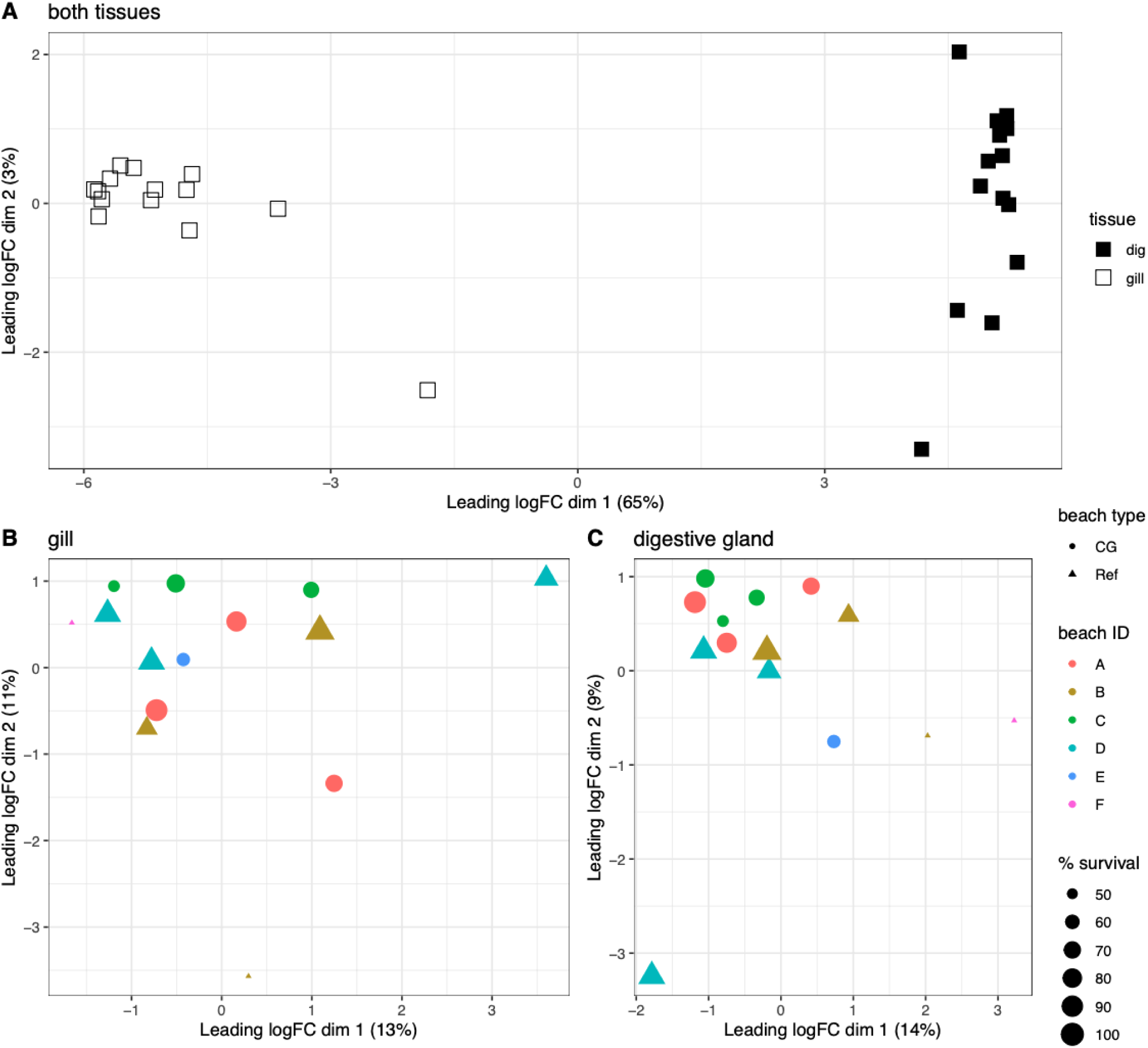
(A) MDS plot for all samples showing clustering based on tissue type (gill or digestive gland; dig) for Pacific littleneck clams sampled from plots P1–P18 on Beaches A−F. MDS plots for (B) gill and (C) digestive gland tissues individually where each datapoint is plotted as an unmaintained clam garden (CG, circles) or reference beach (Ref, triangles), coloured by beach identifier, and sized by the percentage of survival at the beach. The gill sample plot below −3 in dimension 2 is from plot P12, and the digestive gland sample that is below −3 in dimension 2 is from plot P9.

### 3.4 | Tissue-specific expression

Samples from each tissue were then analyzed separately by conducting tissue-specific filtration and normalization. The gill and DG samples had on average (± SD) 12.9 ± 3.0 M and 15.5 ± 2.9 M reads aligning per library and the datasets were found to have 28,391 and 24,699 transcripts expressed, respectively, after filtering for low expression. Inspecting transcripts that were present in only one of the tissue types identified 8,795 (31.0%) gill-specific transcripts and 5,103 (20.7%) DG-specific transcripts (Table 5; *SD* Additional File S2). The highest expressed tissue-specific transcripts are shown in *SI* Figure S6 (N = 200 for each tissue).

**TABLE 5.** Differentially expressed gene overview comparing beach type (clam garden (CG) vs. reference beach (Ref)), beach survival (high vs. medium vs. low), tissue type (gill vs. digestive gland), or expressed in background. Note that here differential expression is considered when *p* ≤ 0.001 and log2 fold-change > 0.58 (1.5-fold overexpressed) or < −0.58 (1.5-fold underexpressed).

| Comparison | Result | # Transcripts in gills | # Transcripts in digestive gland |
| --- | --- | --- | --- |
| **Tissue type (gill vs. digestive gland)** | Gill vs. digestive gland | 8,795 | 5,103 |
| **Beach type (CG vs. Ref)** | Overexpressed CG | 40 | 11 |
|  | Underexpressed CG | 14 | 26 |
| **Beach survival (high vs. medium vs. low)** | Overexpressed high vs. medium | 11 | 4 |
|  | Overexpressed medium vs. low | 14 | 7 |
|  | Overexpressed low vs. medium | 26 | 34 |
|  | Overexpressed medium vs. high | 6 | 18 |
| **Background** | N/A | 28,391 | 24,699 |

Tissue-specific transcripts were compared to all expressed genes in that tissue to identify enriched GO categories in the tissue-specific list. Gill-specific transcripts were enriched for response to stimulus (N = 1,839), signal transduction (N = 1,184), biological adhesion (N = 496), ion transport (N = 494), defense response (N = 402), and G-protein coupled receptor signalling pathways (N = 210) (all *p* < 0.0001), among others (*SD* Additional File S3). DG-specific transcripts were enriched for biological adhesion (N = 308), defense response (N = 263), immune response (N = 223), ion transport (N = 201), innate immune response (N = 149), organonitrogen compound catabolic process (N = 130), defense response to bacteria (N = 82), and humoral immune response (N = 58) (all *p* < 0.0001), among others (*SD* Additional File S3).

### 3.5 | Effect of unmaintained clam gardens on clam gene expression

The gill dataset showed only 13% and 11% of the total variation in an MDS analysis explained by dimensions 1 and 2, respectively (Figure 3B). No clear clustering was observed in the gill samples by beach type (*i.e*., CG vs. Ref) nor by any other evaluated variables (*e.g.*, survival, sand, silt, or carbonates). There were two outlier samples (P12 from beach B and P7 from beach D) in the MDS plot, both of which were Ref beaches and had 45% and 80% survival, respectively.

The DG dataset showed only 14% and 9% of the total variation in an MDS analysis explained by dimensions 1 and 2, respectively (Figure 3C). Some grouping was observed along dimension 2 with CG samples clustering together, except for P15 from beach E. Ref samples were more spread out along both axes. A gradient in survival was observed across dimension 1, with high survival being positioned in negative dimension 1, medium survival near 0-1, and low survival in positive dimension 1, although the trend was not universal for all samples (*e.g.*, P5). Percent carbonate, which was correlated with survival (see above), also trended along dimension 1 in the opposite direction to survival (*i.e.*, high survival being associated with low percent carbonate, *SI* Figure S7).

Differential expression between CG and Ref was evaluated for each tissue separately (Table 5). The gill had 54 transcripts differentially expressed (FC > 1.5, *p* ≤ 0.001, no multiple test correction (MTC)) (see Discussion; *SD* Additional File S2). Of these transcripts, 35 had identifiers recognized by DAVID Bioinformatics, but no Gene Ontology (GO) enrichment was identified (*p* > 0.01). Most of these transcripts were overexpressed in the CG samples (N = 40), including several transcripts associated with immune/apoptotic or stress-related functions, such as two transcripts annotated as *tumour necrosis factor (TNF)*-related (FC > 1.5), one as *death effector domain (DED)*-related (*i.e.*, top overexpressed transcript, FC > 71.5), and *universal stress protein A-like protein* (FC > 4.7). In total, 14 transcripts were underexpressed in the CG samples, including a *heat shock 70 kDa protein 12a* (FC = 17.9). A full list of differentially expressed genes is available in *SD* Additional File S2.

The DG had 37 transcripts differentially expressed between CG and Ref samples. This included 11 overexpressed and 26 underexpressed transcripts in the CG samples (*SD* Additional File S2). Using the full, bidirectional list of differentially expressed transcripts, several enriched GO categories were observed (*SD* Additional File S3), including biological process categories iron ion homeostasis (N = 3, *p* = 0.008) and negative regulation of hydrolase activity (N = 5, *p* < 0.001). The highest overexpressed transcript in the CG samples was *replicase polyprotein* annotated from the *cricket paralysis virus* (FC > 500), which suggests that viral proteins were also captured in the study and that they were differentially abundant between beach types. Multiple complement-, iron-, and protein-folding-related transcripts were observed as differentially expressed in the DG samples, including overexpression of a transcript annotated as *c1q domain* (2.3-fold), *thrombospondin-1* (2-fold), *soma ferritin*, and *cytochrome P450* (FC > 1.5). Transcripts under-expressed in the DG samples in the CG plots included *interferon-induced very large GTPase 1* (FC > 8.5), *apoptosis inhibitor IAP* (4.3-fold), *low affinity immunoglobulin epsilon Fc receptor* (FC > 2), *c-type mannose receptor 2*, and *platelet endothelial aggregation receptor 1* (FC > 1.5), among others (*SD* Additional File S2).

Consistent responses to the CG plots were investigated between the tissues, and two of the 91 unique differentially expressed transcripts were found to be differentially expressed in both tissues. This is notable given that the datasets were normalized, filtered, and analyzed separately. This included the underexpression in the CG plots of a transcript annotated as *von Willebrand factor type D domain (vwd)* and of a transcript annotated as *platelet endothelial aggregation receptor 1* (*pear1*; Figure 4) in both tissues.

**FIGURE 4.**
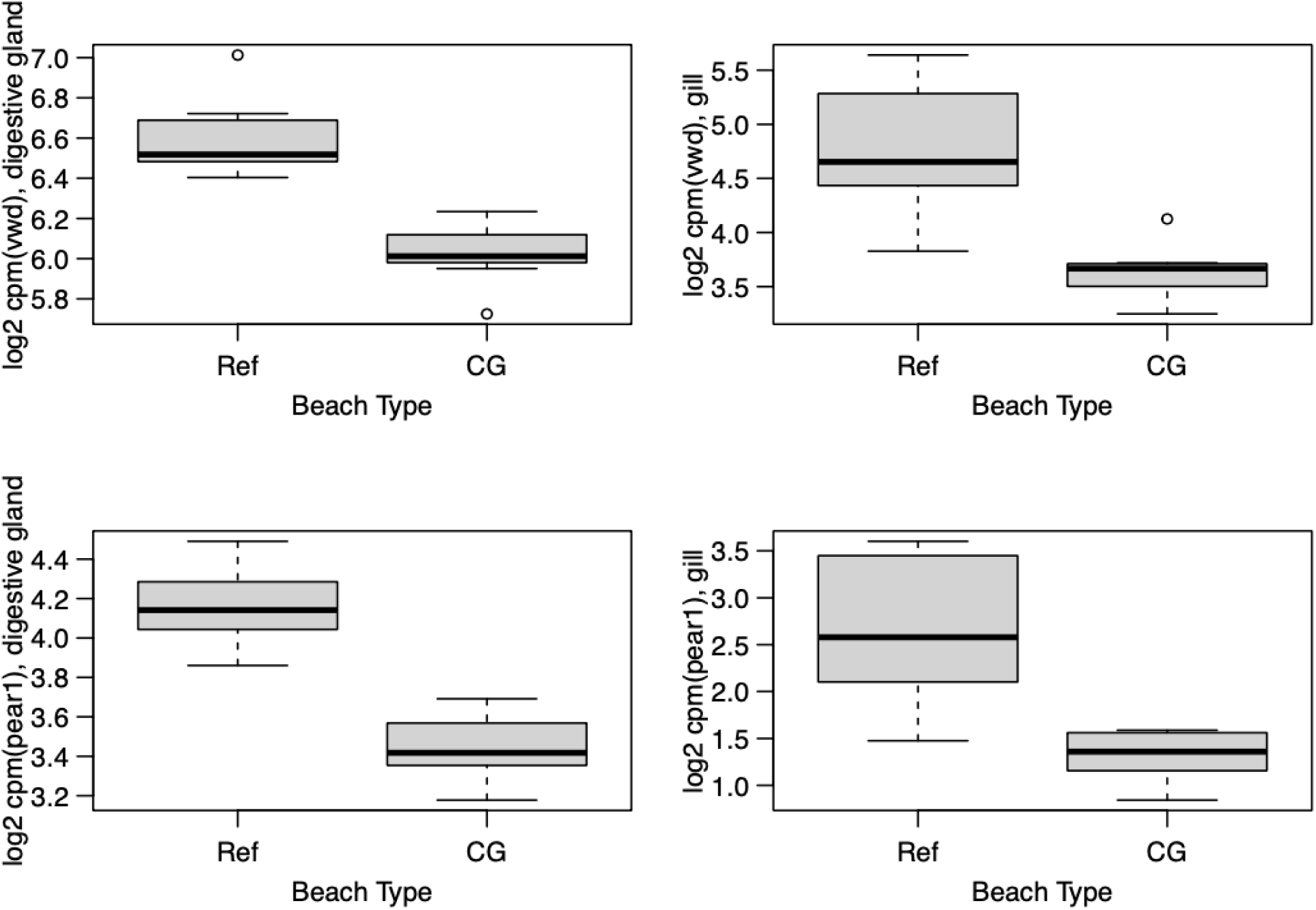
Box plots showing expression of genes of interest identified in both digestive glands and gills of Pacific littleneck clams, which were underexpressed in clam gardens (CG) relative to reference beaches (Ref). vwd = *von Willebrand factor type D domain*; pear1 = *platelet endothelial aggregation receptor 1*.

### 3.6 | Gene expression responses and differential survival

Given the differential survival observed among the beaches and plots, genes correlating with survival were investigated to determine what might be driving these differences and to potentially identify genes that are related to survival or mortality in this non-model species. Genes were of particular interest if they were differentially expressed between low and medium survival, between medium and high survival, or incrementally differentially expressed at each step between high, medium, and low survival (see *Methods* for beach survivorship classification).

The majority of genes significantly associated with survival in the gill were differentially expressed between the low and medium survival groups, rather than between medium and high survival (Table 5). Transcripts overexpressed in the low survival group relative to the medium survival group (N = 26) were involved in functions such as heat-shock, interferon response, and ligase activity (*SD* Additional Files S2, S3). For example, *heat-shock protein (hsp) 90* (FC > 60) and three different transcripts annotated as subunits of *hsp70* (FC > 2) were overexpressed specifically in the low survival group. Transcripts overexpressed in the medium survival group relative to the high survival group included six transcripts of various functions (*SD* Additional File S2). Fewer transcripts were overexpressed in the higher survival group (*i.e.*, high vs. medium, N = 11; medium vs. low, N = 14) and these generally were involved in immune functions, but also various other functions. Overexpression of *toll-like receptor 1* (FC > 2) and *complement C1q-like protein 4* (FC > 3.5) was observed in medium survivors relative to low survivors (Figure 5).

**FIGURE 5.**
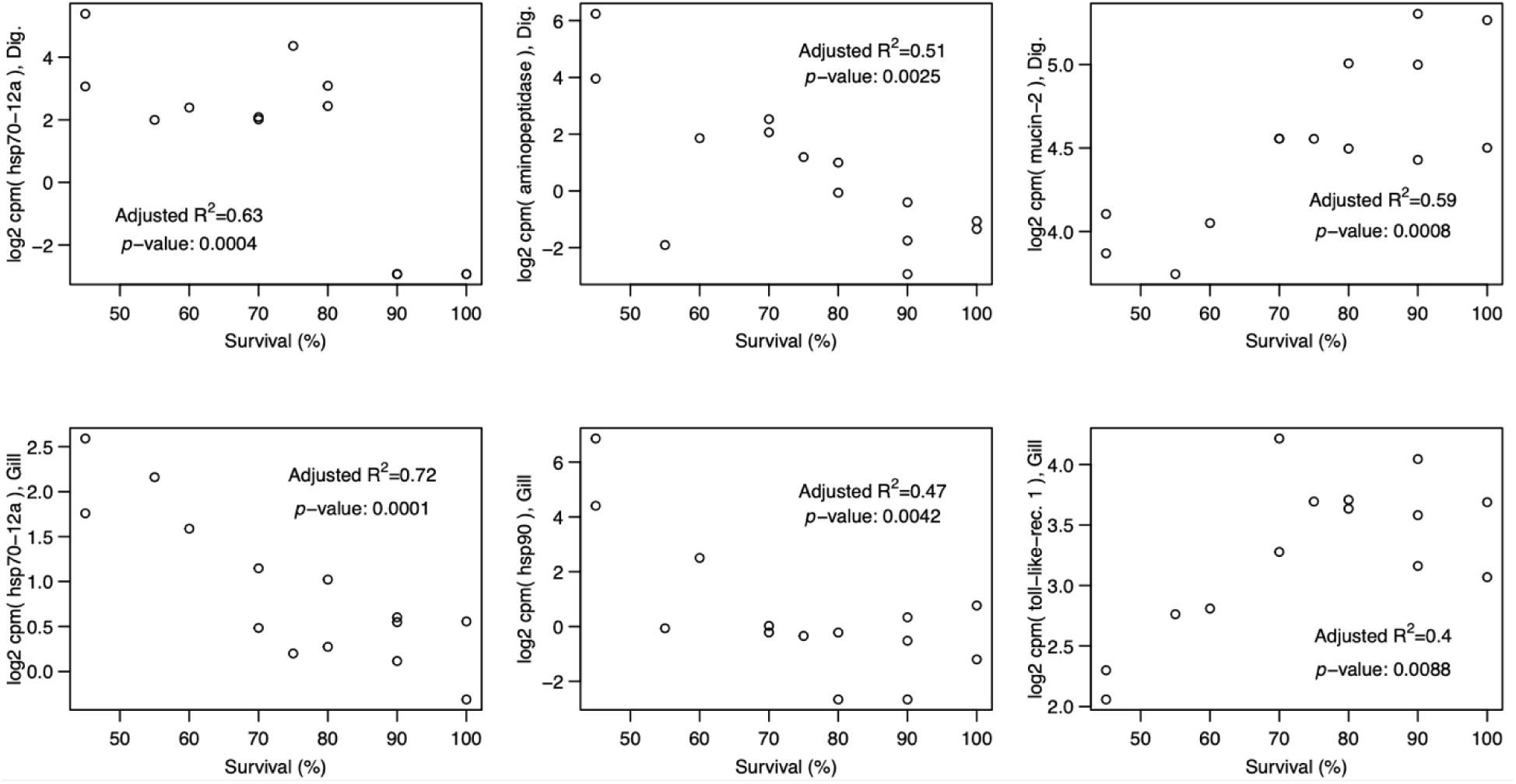
Linear regression of differential expression of selected genes of interest versus percent survival in Pacific littleneck clams. Transcript expression and their correlation to survival include (top left to bottom right) *heat shock protein family A member 12A* (*hsp70-12a*), *aminopeptidase*, and *mucin-2* in the digestive gland (DG) and *hsp70-12a*, *hsp90*, and *toll-like receptor 1* in the gill. Note: adjusted R^2^ and *p-*values presented here are from linear models of log2 counts per million for each transcript against percent survival as a numeric variable.

In the DG, the majority of significant survival-associated transcripts were overexpressed in low relative to medium surviving clams (N = 34) and were generally involved in immune system, heat-shock, or transport functions. Transcripts annotated as *hsp70 12a* and *12b*, as well as several aminopeptidases, were overexpressed in low relative to medium and in medium relative to high survival (shown correlated with survival in Figure 5). All the overexpressed genes in medium relative to high surviving clams (N = 18) were also overexpressed in the low relative to medium survival group comparison. Very few genes were overexpressed in the high survival clams relative to medium (N = 4) or the medium relative to low survival (N = 7, and these transcripts had various functions, including a transcript annotated as *mucin-2 protein* (Figure 5).

## Discussion

### 4.1 | Effects of unmaintained clam gardens on clam and beach characteristics

Results from the present study showed that there was no significant enhancement of growth or survival on unmaintained clam garden (CG) beaches relative to reference (Ref) clam beaches. Using clams of similar sizes that were collected on the same days and location as the present work, Salter (Salter, 2018) also reported no significantly enhanced growth or survival on unmaintained CG compared to reference clam beaches in Kanish and Waiatt Bays. It is important to note that in the present study the clam gardens were unmaintained and therefore were not being tended according to traditional tending practices, which may influence the impact of the garden on clam survival, growth, and transcriptomic profile, as well as various abiotic characteristics (*i.e.*, sediment grain size, carbonate content, organic content). Salter (2018), however, did include some CG traditional tending practices (*i.e.*, adding a 1 cm-thick layer of crushed shell mixture three times throughout the summer) on replicate treatment plots on both beach types and reported a positive effect of the shell hash treatment on growth and survival in both CG and reference clam beaches. Cox et al., reported that even without traditional tending clam gardens contained greater estimated bivalve biomass, distinct infaunal and epifaunal communities and increased topographic complexity compared to reference sites (Cox et al., 2024a; Cox et al., 2024b; Cox et al., 2019). Interestingly, Cox et al., also found that a combination of 45-60% gravel and less than 31% bivalve shells promotes the biologically diverse epifaunal communities observed within clam gardens (Cox et al., 2024a). These results confirm the importance of historical Indigenous practices in maintaining clam gardens and promoting an optimal environment for clam growth and survival, and demonstrate that it is not simply the physical structure of the clam garden wall that creates the ideal clam habitat.

Some trends were observed in the transcriptomic data that suggest that the unmaintained CGs had a transcriptomic effect on the clams, albeit not a strong one. First, the CG and Ref samples showed a slight separation in the PCA of DG tissue, but not gill tissue. Second, although very few genes were differentially expressed, the same differentially expressed genes were consistently identified in both tissues, including transcripts annotated as *von Willebrand factor type D domain (vwd)* and *platelet endothelial aggregation receptor 1* (*pear1*) both of which were downregulated in CG. The *vwd* domain is present in many proteins, including mucins and other extracellular glycoproteins, and one *vwd* transcript in particular was up-regulated in response to heat shock in the Pacific oyster (*Crassostrea gigas*) (Zhang et al., 2015). The function of *pear1* in shellfish remains unknown; in vertebrates it is a cell membrane protein involved in platelet aggregation. Homologous genes for platelet activation are present in the Hong Kong oyster (*Crassostrea hongkongensis*) and are regulated in response to hyposalinity (Xiao et al., 2018). In terms of the other genes differentially expressed by clams in CGs, the general functions were not clear. Immune genes and stress-response genes were found both over- and under-expressed in CGs versus Ref beaches (*e.g.*, a heat-shock protein transcript was found to be underexpressed in the CG and a universal stress protein-annotated transcript was overexpressed in the CG). In general, the functions of these transcripts in the Pacific littleneck clam responding to the CG factor will be useful to continue to characterize as the genomic information develops for this species and additional transcriptomic studies are conducted in relation to environmental stress in maintained or unmaintained clam gardens. These molecular phenotypes may indicate a response to the unmaintained CG from the 16-week *in situ* study period that is below the level of phenotypic growth and survival responses, as these were not found to differ based on beach type. Further work with longer study periods would be required to determine whether these transcriptome signatures precede macro-phenotypic responses.

Sediment grain sizes, percent carbonate content, and percent organic matter did not differ significantly between unmaintained CG and Ref beaches in the present study, although there was significant variation within both beach types among locations. For example, carbonate content varied from 4 to 10% in CGs and from 2 to 15% in Ref beaches, while silt varied from 0 to 4% and from 1 to 5%, respectively. This is somewhat contrary to Salter’s (2018) observations that untreated CGs contained 2.8 – 12.9 times more carbonate and a smaller percentage of silt than non-walled (Ref) beaches (Salter, 2018). The reason for this discrepancy is not known and would be valuable to address with future studies. The lack of differences observed in the present study in growth and survival, and minor effects in transcriptome responses due to unmaintained CGs may be due to the lack of tending of the clam gardens, as is traditionally conducted. Future studies would benefit from fully examining the impacts of tending practices on sediment characteristics alongside clam growth and survival, which would involve clam garden beach restoration and active maintenance for long periods, with further analysis of impacts on shellfish molecular phenotypes and productivity.

### 4.2 | Abiotic variation among beach location and effects on clam growth and survival

Across the six sampling locations, beaches differed in most of the variables examined (*i.e.*, biotic response variables: growth, survival; abiotic characteristics: organic level, carbonate concentration, grain sizes). Beaches clustered based on sediment characteristics, which correlated to beaches in similar geographic or wave-influenced locations, and differential growth and survival were noted among the beach locations. Given these trends, abiotic factors potentially involved in survival as well as molecular responses to survival were characterized.

The sedimentary environment is very important for clam growth and survival (Joo et al., 2021). Littleneck clams typically live as infaunal burrowers between 5 and 15 cm depth, in soft sand or sandy mud in the intertidal zone (Lazo, 2004). In the present study, growth was positively affected by sand, and fine sand, and negatively affected by small rocks, and silt, while survival was positively affected by coarse sand, and sand, and negatively affected by carbonates, organics, small rocks, very fine sand, and silt Joo et al. (2021) found that grain size and sorting of the sediment determined the amount of pore-water dissolved oxygen and organic matter, both affecting growth and survival of juvenile Manila clams, with optimal survival and growth linked to poorly-sorted sediment with an average grain size of medium sand (180 – 335 µm). Similarly, the grain size of the sand in the present study was 250 – 500 µm, and this grainsize had a significant positive effect on survival. A clam tending revitalization project in Burrard Inlet observed that water flow had a positive effect on native clam density and biomass while sediment grain size had little effect (Guttmann, 2022). Salter (2018) found increased water residency time positively affected clam biomass and growth and reported a weak negative effect of silt on clam density and biomass. Contrary to the present study, Salter (2018) also found a positive effect of increased sediment silt content on clam biomass in clam transplant experiments. A multi-year study looking at environmental factors affecting growth in the hard clam *Venus mercenaria* (now *Mercenaria mercenaria*) found a consistent decrease in growth with increasing sediment silt-clay content, with all results showing a negative relationship between growth and fineness of the sediment (Pratt & Campbell, 1956). Permeability of the sediment is an important factor for pumping activity of burrowing filter feeders, which is essential for respiration, excretion, and nutrition and Pratt et al. (1956) found permeability to be an inverse function of the sediment silt-clay content. In addition, commercial Manila clam beds studied in Spain had higher survival of clams planted in sand-gravel than in those placed in mud (Cigarria & Fernandez, 2000). Concurring, the clam *Gomphina veneriformis*, which was buried in sand and silt sediment mixtures, had increased mortality with increasing silt (Li et al., 2021). Likewise, the juvenile bamboo clam (*Solen grandis*) reared in different sediments had, as also seen in the present study, decreased growth and survival rates with increasing fine silt (Chen et al., 2008).

Sediment carbonate content had a significant negative effect on clam survival in the present study, where sediment with more than 8% carbonates had markedly decreased growth and survival. Similarly, Munroe (2016) found that the average daily growth of early post-settled Manila clams over two years was negatively correlated with sediment carbonate content (Munroe, 2016). This is not a universally observed trend in the literature (Groesbeck et al., 2014; Salter, 2018) and counters expectations given the expected importance of carbonate for clam shell growth and early recruitment. The addition of crushed shell hash alters sediment calcium-carbonate saturation states, as it increases surface sediment porewater aragonite content and pH (Green et al., 2009). For instance, there was a three-fold increase in hard clam burrowing and recruitment by adding crushed shell hash to a final concentration of 8% of wet sediment weight (Green et al., 2012). Salter (2018) increased carbonate content significantly (1.9–7.2 times) by addition of shell hash, which increased clam growth and tissue biomass in both clam garden and control beaches after six months (Salter, 2018). Guttman (2022) when examining clam tending practices in Burrard Inlet found that sediment carbonate had a positive effect on native clam density and biomass although when compared to water flow it was imprecise and relatively less important (Guttmann, 2022).

Crushed shell hash also changes the physical structure of the sediment as, like gravel, it can create interstitial spaces in the sediment, thereby increasing pore-water dissolved oxygen, organic-matter content, and substrate stability, as well as providing protection from predators for juvenile clams (Joo et al., 2021; Sponaugle & Lawton, 1990; Thompson, 1995). Thus, there may be multiple benefits of adding shell material for bivalve growth/survival that are not linked to carbonate concentration *per se*. The effect of shell hash or crushed shell on bivalve growth and survival is likely dependent on a wide variety of factors including amount of shell, size and compactness of shell particles, species generating the shell hash, species/size of bivalve grown, stocking densities, tidal height, temperature, and types/densities of predators. For example, the application of very finely crushed shell hash could have a negative effect by decreasing interstitial spaces in the sediment and thus, as seen with silt, decreasing sediment permeability. Recent studies investigating the efficacy of shell hash to mitigate acidification of intertidal sediments found it unsuccessful at raising pore-water pH of acidic sediments (Beal et al., 2020; Doyle & Bendell, 2022). Doyle et al. (2022) found the addition of shell hash to be site dependent as it reduced variation in pH at an intertidal site with pH 8.03, but did not mitigate the pH of mud flats with pH 7.59. More research is required to comprehensively understand the effects of shell hash on bivalve growth and survival.

### 4.3 | Pacific littleneck clam transcriptomics

In non-model organisms, such as the Pacific littleneck clam, advances in molecular ecological and physiological understanding of the species can be advanced through transcriptomic tools and the results these tools generate. Here it was clear that the reference transcriptome generated *de novo* was necessary for the study, given that the closest relative genome (*i.e.*, Manila clam) resulted in a very low alignment rate with the Pacific littleneck clam reads. Further, the use of different tissues in the present study revealed high levels of tissue-specific expression (*i.e.*, 21`-`31% of the transcripts expressed in the tissue). As multiple individuals were included in each sample (*i.e.*, pools), the true number of individuals sampled for the tissue-specific expression analysis is greater than the number of libraries, increasing the reliability of the tissue-specific transcripts. The expression analysis indicated that for both tissues, tissue-specific transcripts were involved in signal transduction, defense and immune response, and ion transport, among other processes. Therefore, although the specific transcripts were different for each tissue, similar functions were enriched in the tissue-specific gene lists for both the gill and the DG, although exceptions were observed.

The presence of differential survival among the locations provided a valuable opportunity for analyzing transcripts that are associated with mortality/survival. These characterizations, over time through multiple transcriptome studies generating more evidence, can begin to characterize specific transcripts that are involved with specific stress types (Evans et al., 2011; Miller et al., 2017; Sutherland et al., 2012). This can enable the development of Ecological Association Ontology data collection for non-model species (Pavey et al., 2012) and in the future may be used to characterize underlying factors associated with observed mortality or morbidity (*e.g.*, abiotic stress compared to bacterial or viral infection). Although the Pacific littleneck clam remains largely uncharacterized at the molecular level, some of the genes identified in the present study correlated with survival, including differential expression in the gill and DG of heat-shock proteins, immune response genes, and other functional categories. Additional transcriptomic studies on the species in different environmental conditions over various temporal and spatial scales should help to improve these categorizations and identify stressor-specific or generalized response genes.

Differential expression was observed in transcripts involved in immune defense. For example, transcripts annotated as complement C1q protein complex or involved in cell adhesion and oxidation-reduction processes had high expression in high surviving beaches and may indicate increased activity of immune defense responses. Complement C1q proteins are an important component of the innate immune system (Zhang et al., 2014) and are upregulated in response to bacterial challenges (McDowell et al., 2014) and salinity stress (Zhao et al., 2012). Cell adhesions are critical for the development, maintenance, and function of multicellular organisms and are essential for invertebrate immunity (Johansson, 1999).

In the present study—potentially due to the relatively low sample size from pooling individuals and/or due to having fewer plots with a sufficient number of survivors at some beaches, and/or due to the high number of expressed transcripts—the impacts of multiple test corrections were severe on the dataset and, therefore, a stringent *p*-value (*p* < 0.001) was applied without a multiple test correction. Associations of genes to CGs or to differential survival should be considered as preliminary observations and further studies evaluating the functions of these transcripts will be beneficial to understand their role in Pacific littleneck clam transcriptomics.

### 4.4 | Viral presence and clam health

The use of RNA-sequencing for the present analysis identified a differentially expressed virus transcript in the CG versus Ref samples, which could have unexpected effects on either gene expression activity or phenotypic responses, as could any other uncharacterized factor influencing clam growth/physiology (*e.g.*, temperature, beach exposure). The top overexpressed transcript in the DG from CGs was a *replicase polyprotein* from the *cricket paralysis virus* (CPV), which is a positive-sense, single-stranded RNA virus from the family Dicistroviridae in the order Picornavirales. Members of the Dicistroviridae family are widely distributed in nature with the broadest host range of all small RNA viruses in invertebrates (Bonning, 2009). As the Dicistroviridae viral sequences isolated in the present study ranged in size from 1479 to 7371 nucleotides, one can deduce that they represent viral mRNA, instead of viral genomes. Some dicistroviruses can be non-pathogenic or cause subtle impacts (reduced longevity and fecundity), while others result in rapid paralysis (Bonning, 2009).

Even though there was transcriptomic evidence of viral presence in all the beaches, most of the Pacific littleneck clams from the medium- and high-survival groups did not exhibit any signs of pathophysiology. Other clams (mostly from the low-survival group), however, were emaciated or watery and had slightly to extremely pale-coloured digestive glands, an indicator of reduced feeding and potentially compromised function and health. Pale digestive glands could also be the result of paramyxean parasites responsible for disease in marine mollusks, with infection being linked to digestive tropism interfering with food adsorption and causing pale-yellowish digestive glands with thin, watery flesh in both mussels and oysters (Alfjorden, 2017; Carella et al., 2011). In the present study, however, there was no significant parasite-derived gene expression observed and histopathology would be needed to confirm any paramyxean parasite infections. It is also worth noting that the single time point of sample collection in the present study and the analysis of only surviving clams may have reduced the likelihood of viewing viral or parasitic gene expression in these samples.

Although very pale digestive glands and emaciation were observed in clams from beaches with lower growth and survival, histopathology or genetic identification would again be required to confirm any viral gene expression linkages to observed impacts on organismal health. Health status is most likely linked to multiple stressors comprised of abiotic (*e.g*., temperature, sediment composition, water residency) and biotic factors (*e.g*., bacteria, viruses, harmful algae). RNA sequencing combined with *de novo* transcriptome assembly revolutionizes virus detection and allows for the screening of a broad range of symptomatic and asymptomatic virus species in a non-discriminatory manner, thereby displaying a broader picture of the environmental factors present (Nagano et al., 2015). The current dataset has yet to be fully characterized for viral sequences and could be mined further at a later date, particularly as more resources are developed for this species.

## Conclusions

The *de novo* transcriptome assembly of the Pacific littleneck clam enabled the investigation of genes differentially expressed based on clam presence in unmaintained clam gardens relative to reference beaches. No abiotic characteristic differences were observed between the two beach types, no phenotypic responses were observed, and all observed transcriptome responses were minor. It is possible that impacts would be greater if the clam gardens were actively tended. However, significant survival differences among beaches independent of clam garden presence/absence, as well as abiotic characteristic differences, indicated several sediment characteristics significantly correlated with growth and survival, including the unexpected result of carbonate content negatively correlated with survival. The survival differences also enabled the detection of gene expression transcripts putatively related to stress response, and along with tissue-specific characterization of the gill and digestive gland, this study advances our molecular understanding of the Pacific littleneck clam, a non-model bivalve species.

## Supporting information

Supplemental Results

Gene Ontology enrichment results

Differentially expressed genes

Abiotic variables with survival and growth

## Acknowledgements

We acknowledge the K’ómoks (Comox) Nation, the Xwémalhkwu (Homalco) Nation, the We Wai Kai (Cape Mudge) Nation, the We Wai Kum (Campbell River) Nation, the Snuneymuxw Nation, the lək□□əŋən (Songhees) Nation, the WSÁNEĆ (Saanich) Nation, and the Wyomilth (Esquimalt) Nation, who have stewarded and protected the territory we worked within since time immemorial. We thank the Hakai Institute, the Tula Foundation, and the Clam Garden Network for facilitating the research and for their continued support. We also thank Caitlin Smith, Morgan Black, Kayla Suhan, Kayla Long, and Kieran Cox for their invaluable help in the field, Dr. Eric Rondeau for his assistance in the laboratory, and Allie Byrne for creating the map of the experimental sites.

## Data Availability

Raw RNA-sequencing data have been uploaded to SRA under BioProject PRJNA818991, BioSamples SAMN26893882–SAMN26893909.

Supplemental materials, including all abiotic and phenotypic site metadata needed for data analysis, transcript annotation, quantified gene expression counts, sample phenotypes:

Reference transcriptome: https://doi.org/10.6084/m9.figshare.19735753

RNA-seq bioinformatics pipeline: https://github.com/bensutherland/Simple_reads_to_counts

RNA-seq analysis and physiological/abiotic analysis pipeline: https://github.com/raapm/ms_clam_garden

## Additional Files

Additional File S1. Abiotic variables with survival and growth linear model *p* and R^2^ values, including and excluding beach D.

Additional File S2. Differentially expressed genes and background gene lists for both tissues in Pacific littleneck clams. Gene lists include clam garden vs. reference, survival groups, and tissue-specific expression.

Additional File S3. Gene Ontology enrichment results for the clam garden analysis, the survival analysis, and the tissue-specific expression.

Supplemental Results provided, including Figures S1−S7.

