## Supplemental Results for "Effects of ancient anthropogenic clam gardens on the growth, survival, and transcriptome of Pacific littleneck clams (*Leukoma staminea*)"

**Table of Contents:**

| **Figure S1.** Per beach survival and growth | **Page 2** |
| --- | --- |
| **Figure S2.** Per beach starting weight and height | **Page 2** |
| **Figure S3.** PCA of physical and biotic characteristics, without beach D. | **Page 3** |
| **Figure S4.** Correlation plot of physical and biotic characteristics, all beaches. | **Page 4** |
| **Figure S5.** Correlation plot of physical and biotic characteristics, without beach D. | **Page 4** |
| **Figure S6.** Heatmap of tissue-specific expression transcripts showing highest expressed. | **Page 5** |
| **Figure S7.** Percent survival by carbonate content. | **Page 6** |

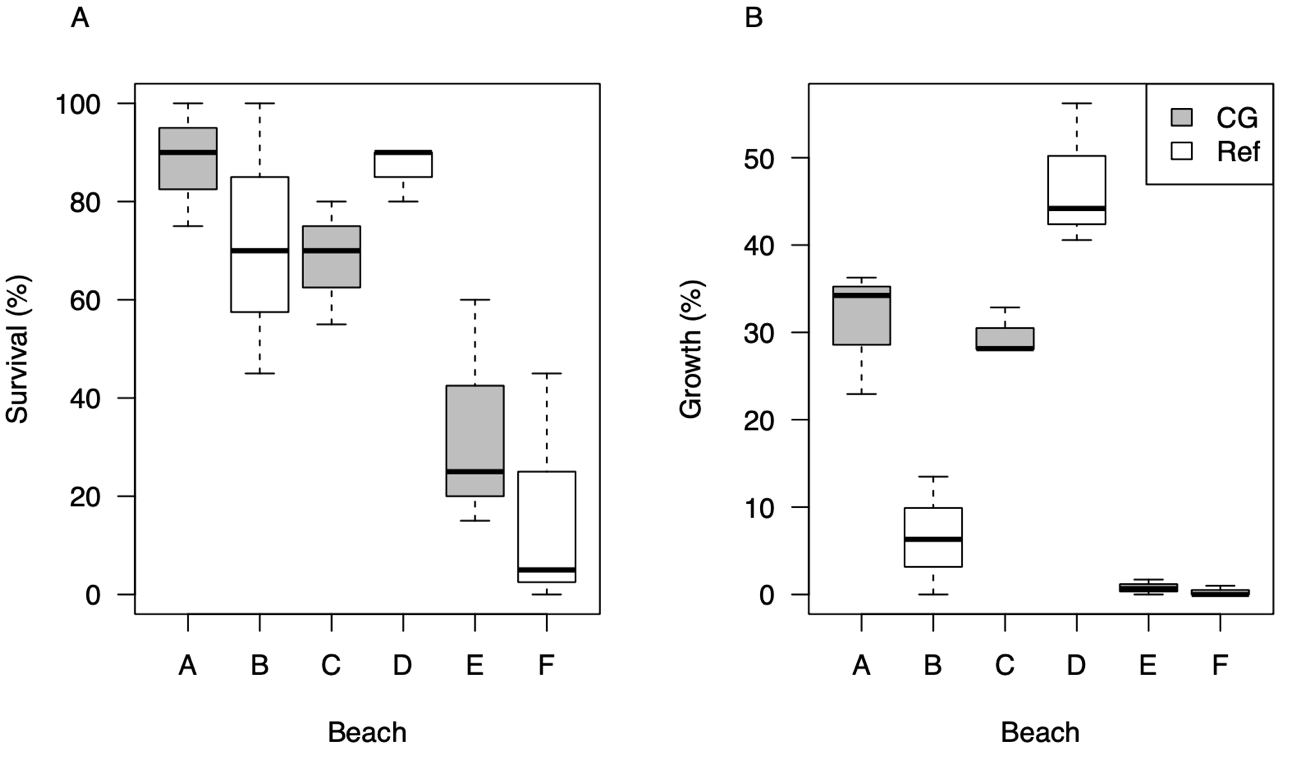

**FIGURE S1.** Boxplots of (A) percent survival and (B) percent growth of Pacific littleneck clams by beach (beaches A–F), with shading denoting whether the location is an unmaintained clam garden (CG) or reference clam beach (Ref).

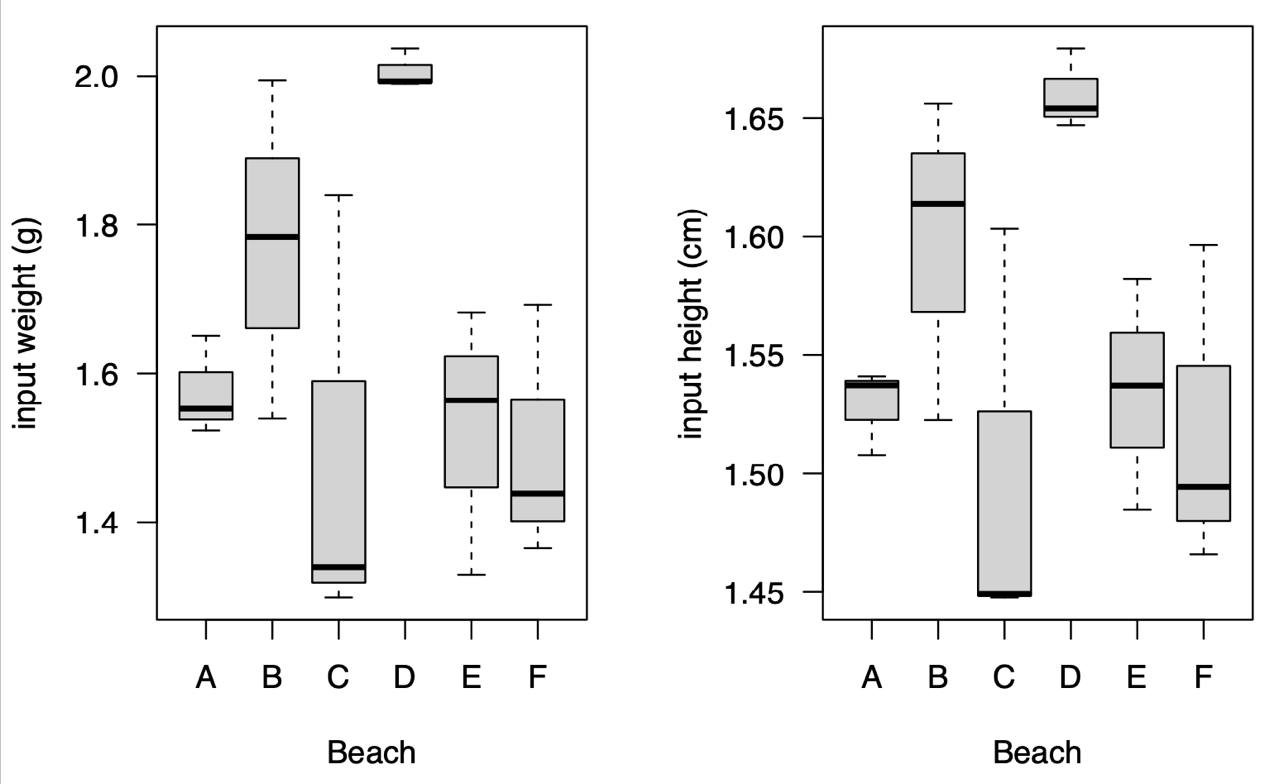

**FIGURE S2.** Boxplots of (A) starting weight in grams and (B) starting height in cm of Pacific littleneck clams by beach (A–F).

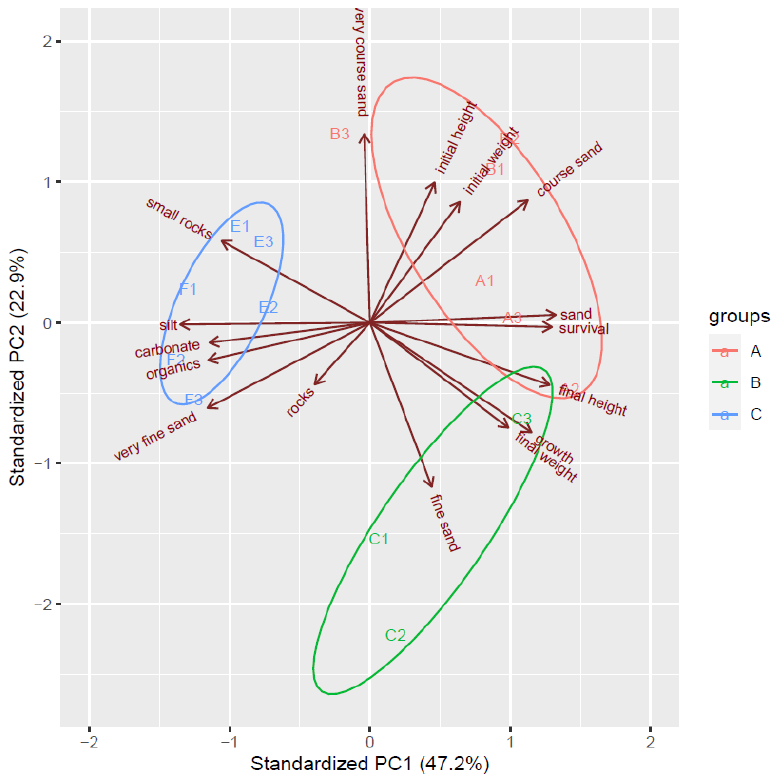

**FIGURE S3.** Principal component analysis biplot based on abiotic and biotic phenotypes showing beach groupings and correlations with beach D excluded (*i.e.*, the beach with an unplanned size difference at experiment initiation). Each beach is denoted as A through F as per Figure 1 locations, and each replicate plot within the beach as 1−3. Colours of plots and ellipses are for visualization purposes only.

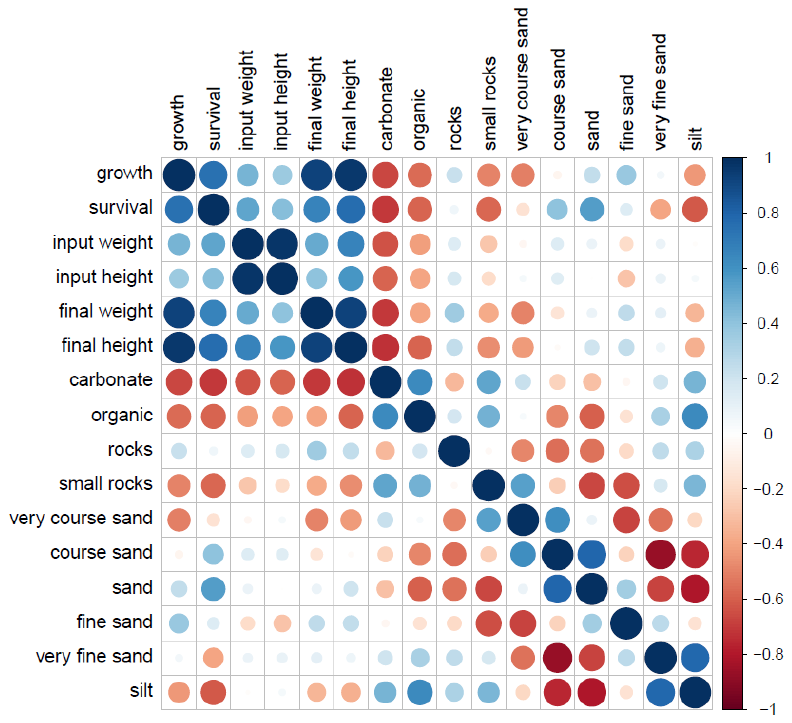

**FIGURE S4.** Pearson correlations between biotic and abiotic variables in the study. Beach D, the beach with unexpected size differences at the start of the study was included in this correlation analysis.

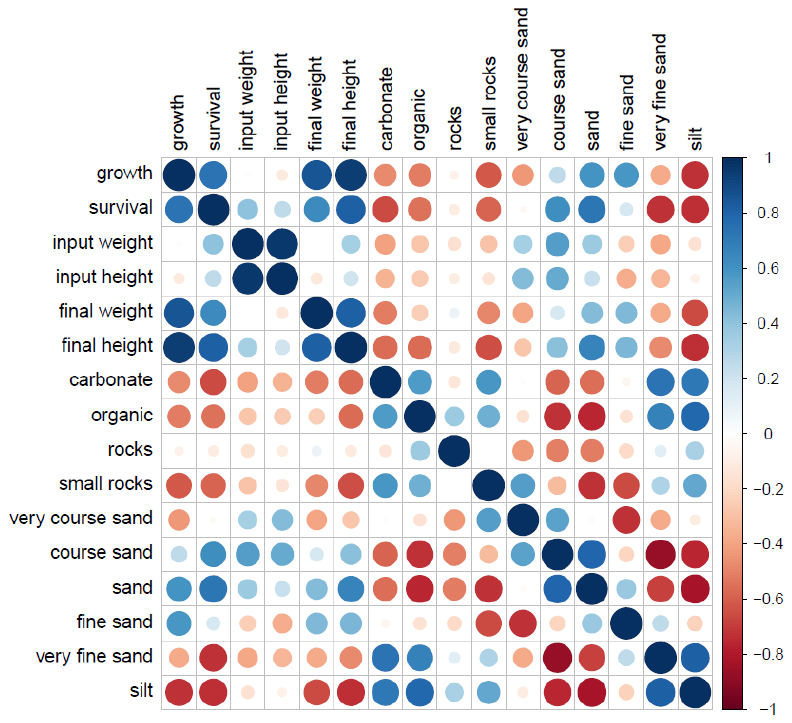

**FIGURE S5.** Pearson correlations between biotic and abiotic variables in the study. Beach D, the beach with an unexpected size difference at the start of the study was not included in this correlation analysis.

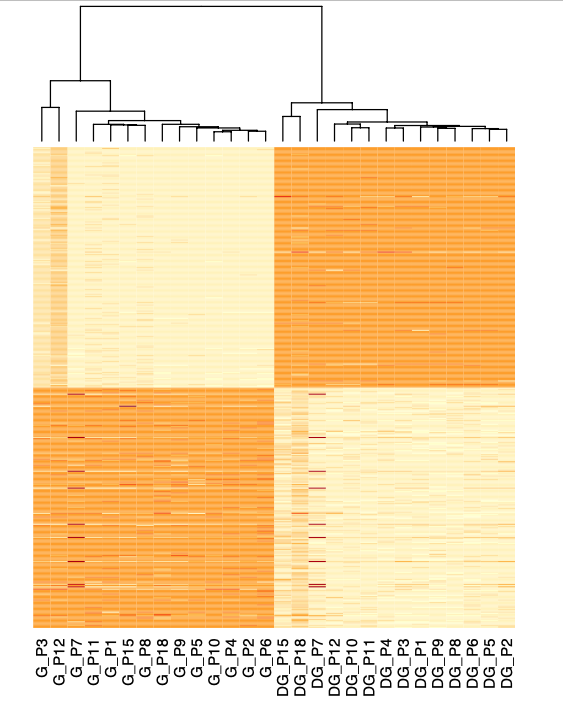

**FIGURE S6.** Tissue-specific expression shown based on the highest expressed tissue-specific transcripts in the gill (G) and digestive gland (DG) for pooled samples within each plot (P1–P18). Samples were clustered based on expression of these transcripts, grouping similarly expressing samples. The higher the expression, the darker the colour of the cell, and each row is an individual transcript. Plot numbers correspond to beaches as follows: Beach A: P1–P3; Beach B: P10–P12; Beach C: P4–P6; Beach D: P7–P9; Beach E: P15; Beach F: P18.

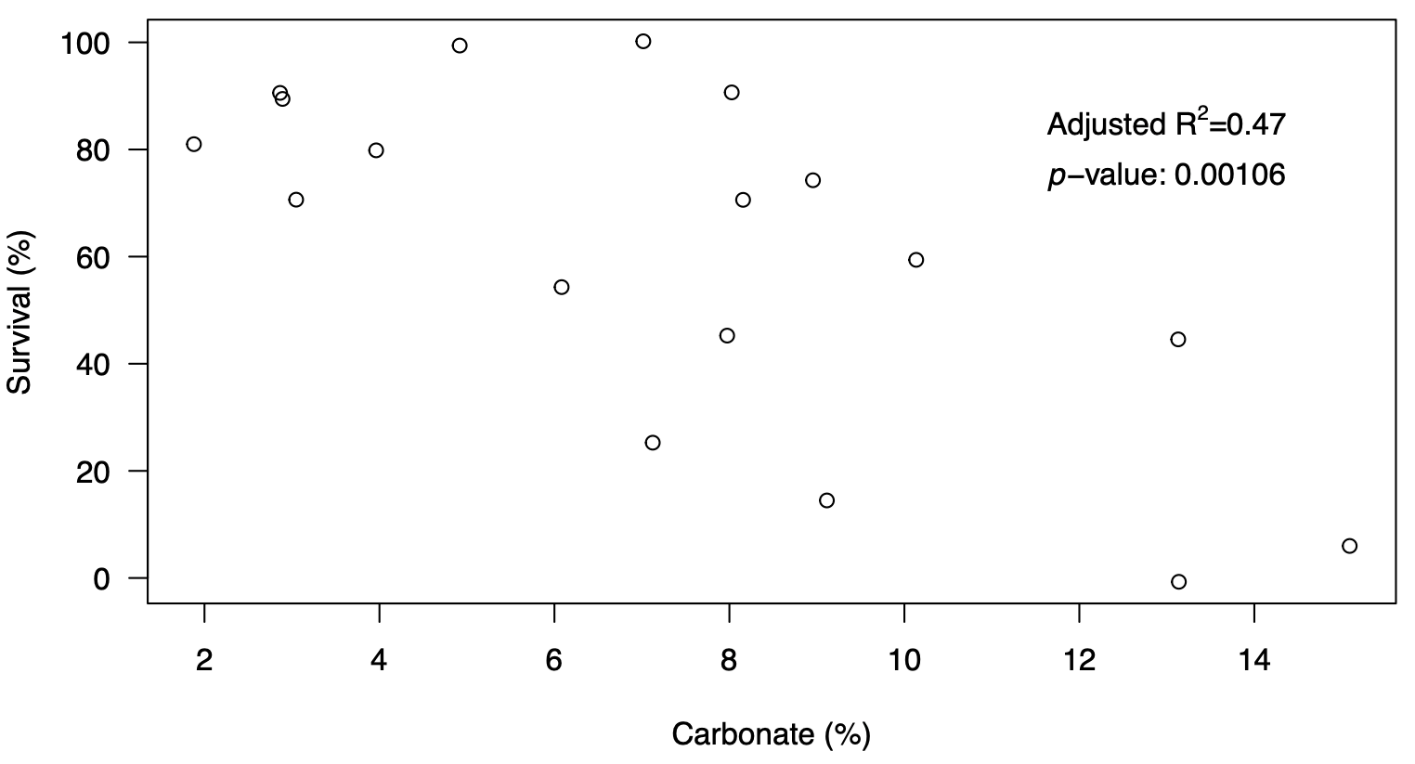

**FIGURE S7.** Linear regression of percent survival of Pacific littleneck clams versus percent carbonate content in sediments. Beach D, the beach with an unplanned size difference at the start of the study, is included in this figure, and when it is not included, the adjusted R^2^ = 0.39 (*p* = 0.008); see *SD* Additional File S1. Points are included with a jitter to avoid overlap. Percent carbonate values are given to the nearest percent and survival to the nearest 5%.
